# High-performance GPCR optogenetics based on molecular properties of animal opsins, MosOpn3 and LamPP

**DOI:** 10.1101/2022.02.07.479375

**Authors:** Mitsumasa Koyanagi, Baoguo Shen, Takashi Nagata, Lanfang Sun, Seiji Wada, Satomi Kamimura, Eriko Kage-Nakadai, Akihisa Terakita

## Abstract

Optogenetics for GPCR signaling is highly valuable but still requires effective and versatile tools with performance evaluation from molecular properties. Here we investigated performance of two animal opsins, mosquito Opn3 (MosOpn3) and lamprey parapinopsin (LamPP) in optical manipulation *in vivo* by using *C. elegans*. MosOpn3 introduced in a nociceptor neurons induced avoidance responses light-dependently with a retinal isomer ubiquitously present in every tissue, like ChR2 and unlike canonical vertebrate opsins. Remarkably, the sensitivity is ~7000 times higher than the case of ChR2 in the light-induced behavior. LamPP introduced in motor neurons induced violet light-dependent stop and green light-dependent go, demonstrating color-dependent manipulation of behaviors using LamPP. Furthermore, our molecular engineering extended the usability of MosOpn3 and LamPP to different signaling cascades and kinetics. Current findings demonstrated that the availability of two animal opsins is equivalent to that of ChR2 in terms of retinal requirement, providing solid strategies for GPCR optogenetics.

## Introduction

G protein-coupled receptors (GPCRs) are transmembrane receptors, which are involved in various cellular and physiological functions including neural responses, cell metabolisms, and hormonal responses^1,2^. GPCRs generally bind a variety of chemical ligands such as odorants, hormones and neurotransmitters, and the extracellular signals are transduced into intracellular signaling (GPCR signaling) *via* heterotrimeric G proteins. The GPCR signaling varies mainly depending on the subtype of G protein alpha subunit and includes cAMP-signaling up- and down-regulated by Gs- and Gi-type G proteins, respectively, and phosphoinositol-signaling for Ca^2+^ elevation mediated by Gq-type G protein. Most animals have hundreds of GPCR genes and human, for example, has ~800 GPCR genes, indicating the importance of GPCR signaling for biological activities^3^. Although structure-function relationships of many GPCRs have been well investigated so far, for comprehensive understanding of GPCR-based physiologies as well as controlling them precisely, optical manipulation of GPCR signaling would be one of the ultimate approaches because of high temporal resolution of light stimulus.

Animal rhodopsins (opsin-based pigments), which underlie vision and non-visual functions such as circadian photoentrainment in varied animals, consist of a protein moiety, opsin, and 11-*cis* retinal as a chromophore in many cases and basically serve as light-sensitive GPCRs^4^. Therefore, opsins have been considered promising tools for optical manipulation of GPCR signaling, and indeed, such optogenetic application has succeeded to some extent using vertebrate visual opsins^5–7^. However, the fact that only a small number of papers on optogenetic research using animal opsins have been published until now compared to those using microbial rhodopsins such as ChR2 for optical manipulation of neural activities suggests that there is still room for improvement. The point would be related to molecular properties of vertebrate visual opsins; 1) they basically form photopigments by binding to 11-*cis* retinal, which is abundant in photoreceptor tissues like eyes, but not in other tissues (11-*cis*-retinal-requirement), and 2) after light absorption their photopigments immediately release chromophore (bleach) to be functionless (bleaching property), both of which are unfavorable characteristics for high performance optical manipulation of GPCR signaling, especially *in vivo*^4^. To overcome the problems associated with the molecular properties, utilization of animal opsins that is more suitable for optogenetic tools should be required.

To date, thousands of opsins have been identified from a wide range of animals from cnidarians to vertebrates, and they are phylogenetically divided into eight or more groups, which is almost consistent with the classification based on G protein selectivity and activation manner^4,8^. With respect to the photochemical property, opsins are basically classified into two types, bleaching opsins like vertebrate visual opsins and bleach-resistant or bistable opsins, which convert to stable active states upon light absorption and in many cases, active states revert to the original inactive state by subsequence light absorption^4,9^, like invertebrate visual opsins^10–13^. The bistable nature appears to be suitable for sustained optical manipulation of GPCR signaling. In fact, a bistable opsin melanopsin (OPN4) was applied to optical control of some physiologies including restoration of vision^14–16^. In addition to melanopsin, we have identified many kinds of bleach-resistant/bistable opsins from both invertebrates and vertebrates^10,17–23^. Among them, we particularly focused on optogenetic potentials of Opn3 and parapinopsin because of their interesting molecular properties, which are recently receiving increasing attention^24^.

Opn3 was first identified from mammalian brain and therefore originally called encephalopsin^25^. Then its homologs were identified from many animals and revealed to be expressed in their various tissues including brain, suggesting their involvement in photoreception in “non-photoreceptive” tissues. We have previously succeeded in functional analyses of members of the Opn3 group^22,26^ and found that one of the members, mosquito Opn3 has a unique property; it forms a photopigment that light-dependently activate Gi and Go-type G protein, when bound to 13-*cis* retinal as well as 11-*cis* retinal^22,27^. Since 13-*cis* retinal is thermally equilibrated with all-*trans* retinal, a retinal isomer ubiquitously present in animals, we have proposed the idea that the Opn3 is applicable to anywhere in the body as an optogenetic tool. In fact, mammalian cultured cells expressing the mosquito Opn3 exhibited light-induced decrease of intracellular cAMP level probably mediated by the Gi activation with all-*trans* retinal addition or even without retinal addition, under the presence of only a small amount of retinoid in serum^22^. The idea was also supported by the recent report showing optogenetic silencing of neurotransmitter release with the mosquito Opn3 *in vitro* and *in vivo*^28^.

Another promising bistable opsin parapinopsin was first identified from catfish pineal and parapineal organs^29^ and thereafter from many lower vertebrate pineal-related organs^17,23,30,31^. Spectroscopic and biochemical analyses revealed that the lamprey parapinopsin is a UV-sensitive bistable opsin, which activates transducin and Gi-type G protein upon UV-light absorption^32–34^. Notably, parapinopsins convert to the stable active state having an absorption maximum at ~500 nm, in green region, which is largely distinct from that of the inactive dark state^32^. The large spectral difference between the inactive and active states allows selective illumination of active state, resulting in its complete recovery to the inactive state^17^. The same is true for the signal transduction level. The G protein activation by parapinopsin were up- and down-regulated by UV and green light illumination, respectively *in vitro* and *in vivo*^33–38^, demonstrating its optogenetic potential for color-dependent on and off of GPCR signaling.

Here, to evaluate performances of the mosquito Opn3 (MosOpn3) and lamprey parapinopsin (LamPP) in optical manipulation of GPCR signaling *in vivo* based on their molecular properties, we focused on *Caenorhabditis elegans*, in which relationships between GPCR signaling and behaviors have been well defined. Importantly, in the case of optogenetics in *C. elegans*, isomeric forms of chromophore retinal can be controlled by exogenously adding specific retinal isomers^6,39^, which is an irreplaceable advantage in testing the chromophore requirement of opsin for functioning *in vivo*, in 11-*cis* form poor condition. In this paper, we showed that MosOpn3 functions *in vivo* under the absence of 11-*cis* retinal with much higher sensitivity compared to ChR2, which is the most used optogenetic tool. We also succeeded in color-dependent manipulation of a *C. elegans* behavior by using LamPP. Together with our demonstration of introducing G protein selectivity of particular GPCRs into MosOpn3 and LamPP, current findings provide versatile and powerful optogenetic tools for manipulating GPCR signaling and various physiologies based on the molecular properties of the two opsins.

## Results

### Optical manipulation of *C. elegans* behavior using the mosquito Opn3

We evaluated the performance of mosquito Opn3 (MosOpn3) for its molecular property-based optical manipulation of GPCR signaling *in vivo* using *C. elegans*. Since MosOpn3 is a Gi/o-coupled opsin^22^, we focused on ASH neurons, a kind of nociceptors, in which chemoreceptors trigger Gi/o-like G protein (ODR-3)-mediated signal transduction upon ligand binding to induce avoidance behavior of *C. elegans*^40,41^ (Fig. 1a). MosOpn3 was introduced into ASH neurons under the control of the promoter of sra-6, a chemosensory receptor mainly expressed in ASH neurons^42^ (Fig. 1b). We obtained several lines of transgenic (Tg) worms that express MosOpn3 in ASH neurons (MosOpn3-worms) with the aid of the pharynx expression of mCherry introduced as a selection marker together with MosOpn3. The expression of MosOpn3 in ASH neurons were confirmed by the expression of GFP, which was designed to be bicistronically expressed with MosOpn3 in ASH neurons. We performed behavioral experiments (Fig. 1c) for a light-induced avoidance of MosOpn3-worms that were fed 11-*cis* retinal-containing *Escherichia coli* (MosOpn3/11-worms). As results, MosOpn3/11-worms exhibited clear avoidance responses by illumination of white light (Fig. 2a upper panels, Supplementary Movie 1a). On the other hand, MosOpn3-worms without a supply of retinal (MosOpn3/NoRet-worms) did not exhibit the light-induced avoidance responses at all (Supplementary Movie 1b), which is consistent with previous observations that the addition of retinal is required for functioning of rhodopsins in *C. elegans*^6,39^. The retinal requirement demonstrated that the light-induced avoidance responses of MosOpn3-worms were not caused by the endogenous light sensor protein, lite-1^43,44^, but indeed by MosOpn3 provably through ODR-3-mediated signaling. Collectively, the results ensure the validity of our experimental conditions including light intensity for investigating the functionality of heterologously expressed animal opsins in *C. elegans*. Importantly, when MosOpn3-worms were fed all-*trans* retinal (MosOpn3/AT-worms), they also exhibited the light-induced avoidance responses (Fig. 2a lower panels, Supplementary Movie 1c) like the case of MosOpn3/11-worms, which can be explained by the unique molecular property of MosOpn3, the pigment formation ability with 13-*cis* retinal thermally generated from all-*trans* form as observed in our previous *in vitro* experiment^22^.

**Fig. 1.**
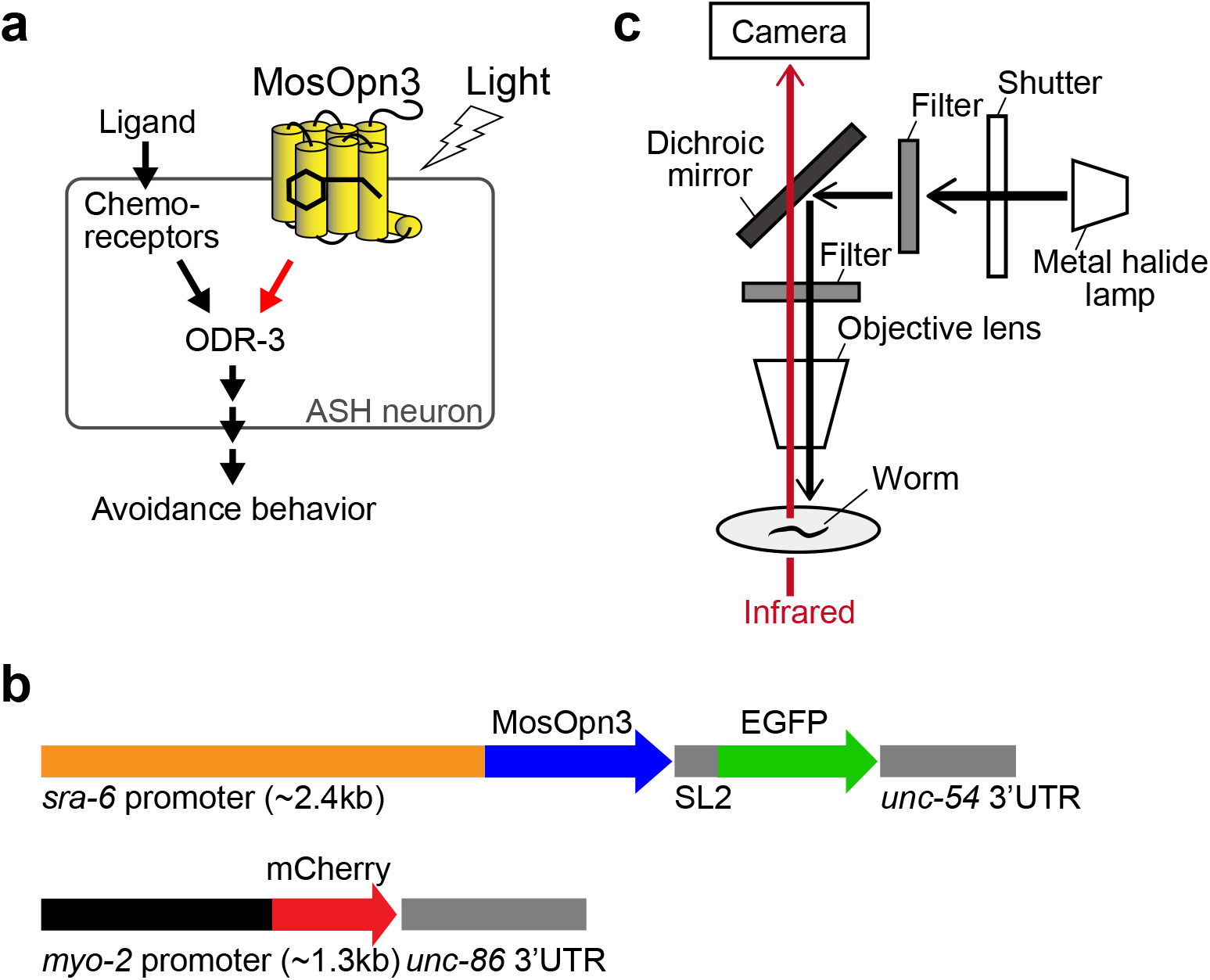
Examination of the functionality of MosOpn3 in ASH neurons of *C. elegans*. **a** The construction for introducing MosOpn3 with EGFP under the *sra-6* promoter. A selection marker mCherry was also introduced under the promotor of *myo-2*. **b** GPCR signaling in ASH neurons. Activation of ODR-3, a Gi-like protein eventually causes avoidance behavior of *C. elegans*. **c** Experimental setup. Worms were continuously monitored with infrared light. Light stimuli were supplied using a dichroic mirror and the intensities were regulated with ND filters.

**Fig. 2.**
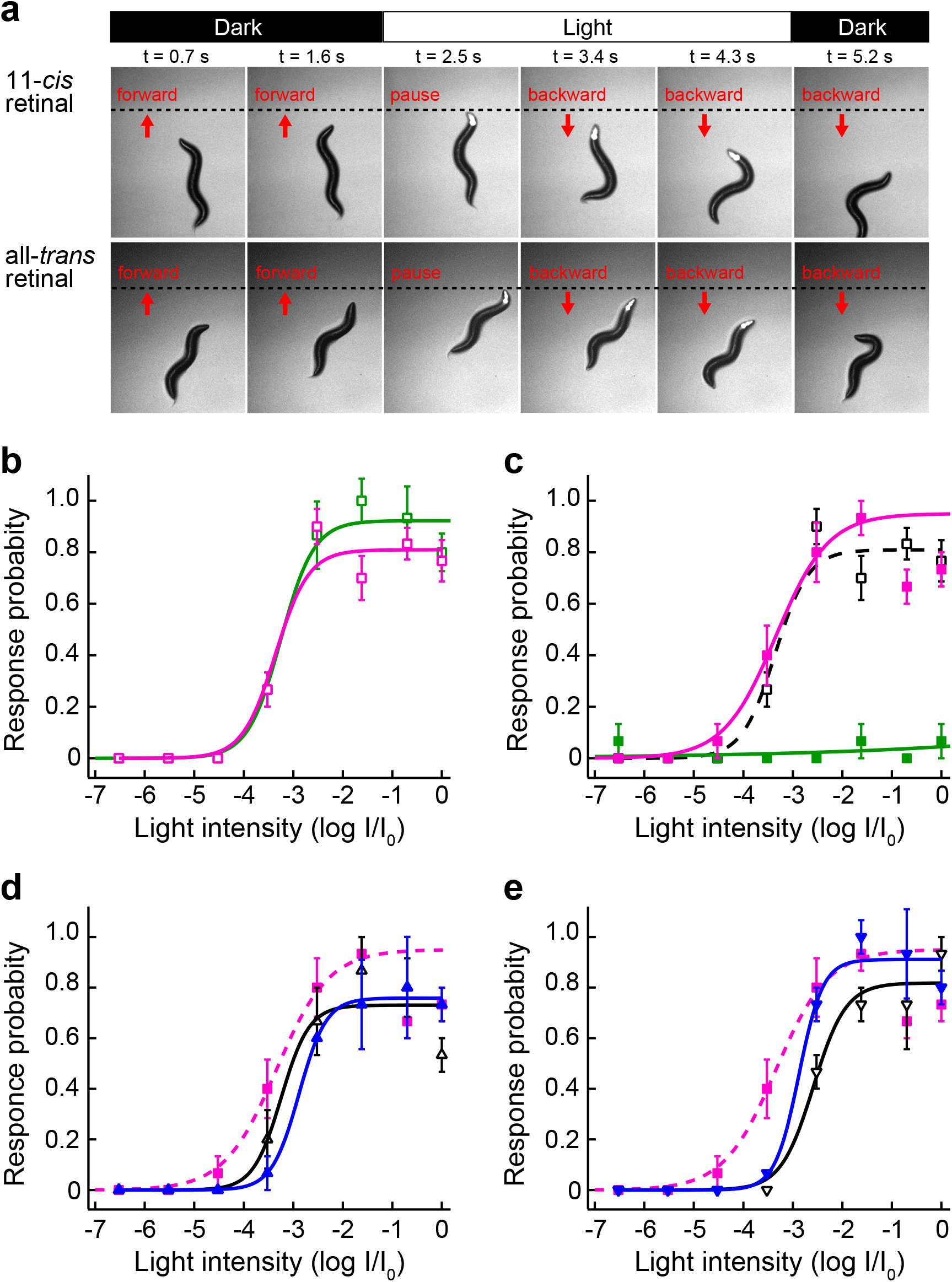
Functionality of MosOpn3 in ASH neurons of *C. elegans*. **a** Snapshot images at 0.7, 1.6, 2.5, 3.4, 4.3 and 5.2 s after recording start, showing light-induced avoidance responses of MosOpn3-expressing Tg worms in the presence of 11-*cis* (upper) and all-*trans* (lower) retinal. The horizontal bar and mCherry signals emitted by the light illumination indicate the timing of white light stimulus. Dotted lines indicated the position of the frontal tip of worms when they stopped moving forward. **b, c** White light intensity-response probability relationships for MosOpn3-worms (magenta squares and curve) and BovRh-worms (green squares and curve) in the minimum necessary amount 11-*cis* retinal (**b**, n=6) and all-*trans* retinal (**c**, n=3). Both Tg worms exhibited a similar light sensitivity in the case of 11-*cis* retinal whereas only MosOpn3-worms responded to light under the presence of all-*trans* retinal. The light intensity-response probability relationship for MosOpn3/11-worms is also indicated in (**c**) for comparison (black open squares and dashed curve), showing a similar performance of MosOpn3/11- and MosOpn3/AT-worms in the light-induced avoidance behavior. **d, e** White light intensity-response probability relationships for MosOpn3/11-worms (black triangles) and MosOpn3/AT-worms (blue triangles) under the presence of 1/1000 (**d**, n=3) and 1/10000 (**e**, n=3) amount of the standard amount of retinal. The data for MosOpn3/AT-worms under the minimum necessary amount retinal were also indicated for comparison (magenta filled squares and dashed curve). I_0_ = ~0.8 mW/mm^2^ (**b**-**e**).

### Comparison of the functionality and efficiency *in vivo* between MosOpn3 and BovRh

To evaluate the advantage of MosOpn3 in optical manipulation of GPCR signaling *in vivo*, we also investigated the light-induced avoidance behaviors of Tg *C. elegans* expressing a bleaching opsin, bovine rhodopsin (BovRh), which activates Gi/o type G protein^45^ like MosOpn3 but does not bind 13-*cis* retinal or all-*trans* retinal unlike MosOpn3. Tg worms established to express BovRh in ASH neurons (BovRh-worms) exhibited the avoidance behavior by white light illumination when they were fed 11-*cis* retinal-containing *E. coli* (Supplementary Movie 1d, e). On the other hand, BovRh-worms did not respond to light when they were fed all-*trans* retinal-containing *E. coli* (Supplementary Movie 1f), which is different from the case of MosOpn3/AT-worms, showing good agreement with molecular properties of respective opsins. We then compared the performances of these two animal opsins in optical manipulation of GPCR signaling *in vivo* quantitatively. We investigated relationships between light intensity and light-induced avoidance response probabilities of MosOpn3- and BovRh-worms with various amounts of 11-*cis* retinal to find the necessary amount of 11-*cis* retinal for functioning in ASH neurons to avoid side effects caused by an excess amount of retinal. The light intensity-response probability relationships revealed that MosOpn3-worms exhibited a similar light sensitivity even when the amount of 11-*cis* retinal was reduced to 1/100, and the sensitivity was slightly decreased under 1/1000 amount of 11-*cis* retinal (Supplementary Fig. 1a). In the case of BovRh-worms, the light sensitivity was also similar even when the amount of 11-*cis* retinal was reduced to 1/100, but the sensitivity was largely decreased under 1/1000 amount of 11-*cis* retinal (Supplementary Fig. 1b). These results indicate that the 1/100 amount is necessary to induce maximum performance for both opsins in this behavior. We then compared performances of MosOpn3- and BovRh-worms in the presence of the minimum necessary amount of retinal. We found that light sensitivities of MosOpn3/11- and BovRh/11-worms are similar, which suggests a similarity in performance of 11-*cis* binding-MosOpn3 and BovRh in the ASH neurons (Fig. 2b). On the other hand, in the same amount of all-*trans* retinal, MosOpn3/AT-worms exhibited the avoidance responses by illuminations of 1/10000~1/1000 of the maximum intensity (I_0_) of light, whereas BovRh/AT-worms did not exhibit any avoidance responses even by illuminations of the maximum intensity of light (Fig. 2c), demonstrating that the sensitivity of MosOpn3/AT-worms is more than 1000 times higher than that of BovRh/AT-worms. Remarkably, the sensitivity of MosOpn3/AT-worms is comparable to that MosOpn3/11-worms (Fig. 2c). In addition, when the amounts of 11-*cis* and all-*trans* retinal added to worms were reduced to 100 times lower level, the sensitivities of MosOpn3-worms under the presence of 11-*cis* and all-*trans* retinal decreased similarly (Fig. 2d, e). Since the decreases of sensitivity are explained by those of formed photopigment amount in *C. elegans*, the similarities in sensitivity between MosOpn3/11- and MosOpn3/AT-worms under varied retinal amounts suggest that MosOpn3 bound to 11-*cis* retinal and MosOpn3 bound to 13-*cis* retinal, which is thermally generated from all-*trans* retinal, can function with similar efficiency in ASH neurons. These results clearly demonstrated that MosOpn3 performs efficiently in ASH neurons to evoke avoidance responses even in the presence of all-*trans* retinal like microbial rhodopsins.

### Engineering MosOpn3 for light-dependent upregulation of intracellular cAMP and Ca^2+^ levels

MosOpn3 forms a photopigment by binding 13-*cis* retinal to activate Gi and Go type G protein in a light dependent manner^22^, which leads to decrease of cAMP. To expand the scope of optogenetic application of MosOpn3, we engineered MosOpn3 to upregulate cAMP and Ca^2+^ levels, through Gs- or Gq-type G protein, respectively by exchanging cytoplasmic region(s) including the third cytoplasmic region, the major determinant of G protein selectivity for Class A GPCRs^45–51^. We chose the ß2-adrenergic receptor (ß2AR) and α1-adrenergic receptor (α1AR), which are generally considered to selectively activate Gs- and Gq-type G protein, respectively, as donor GPCRs. We generated a series of MosOpn3-ß2AR and MosOpn3-α1AR chimeras, in which cytoplasmic region(s) were replaced with those of ß2AR or α1AR according to previous reports^5,50^, and investigated functional conversion. In the case of MosOpn3-ß2AR chimeras, we measured light-induced increases of cAMP level in cultured cells expressing each chimera using the GloSensor cAMP assay, and revealed that all chimeras induced obvious cAMP increases light dependently in contrast to the cAMP decrease in the wild type (WT) even without addition of retinal (Fig. 3a), indicating successful conversion of cAMP regulation from down- to up-regulation by MosOpn3 bound to endogenous retinal in the culture medium. Any MosOpn3-ß2AR chimeras exhibited larger cAMP increases upon light absorption than the previously reported MosOpn3 chimera containing the third cytoplasmic region of jellyfish Gs-coupled opsin^20,26^. Among all, the chimera in which all cytoplasmic regions were replaced with those of ß2AR (MosOpn3-ß2ARiL123C) exhibited the largest cAMP increase and the chimera containing the second, third and C-terminal region of ß2AR (MosOpn3-ß2ARiL23C) was comparable. An interesting finding is that introducing the first cytoplasmic region of ß2AR accelerates the reversion of increased cAMP levels to the basal level, which provides choices of different kinetics of cAMP changes based on the chimera species (Fig. 3a).

**Fig. 3.**
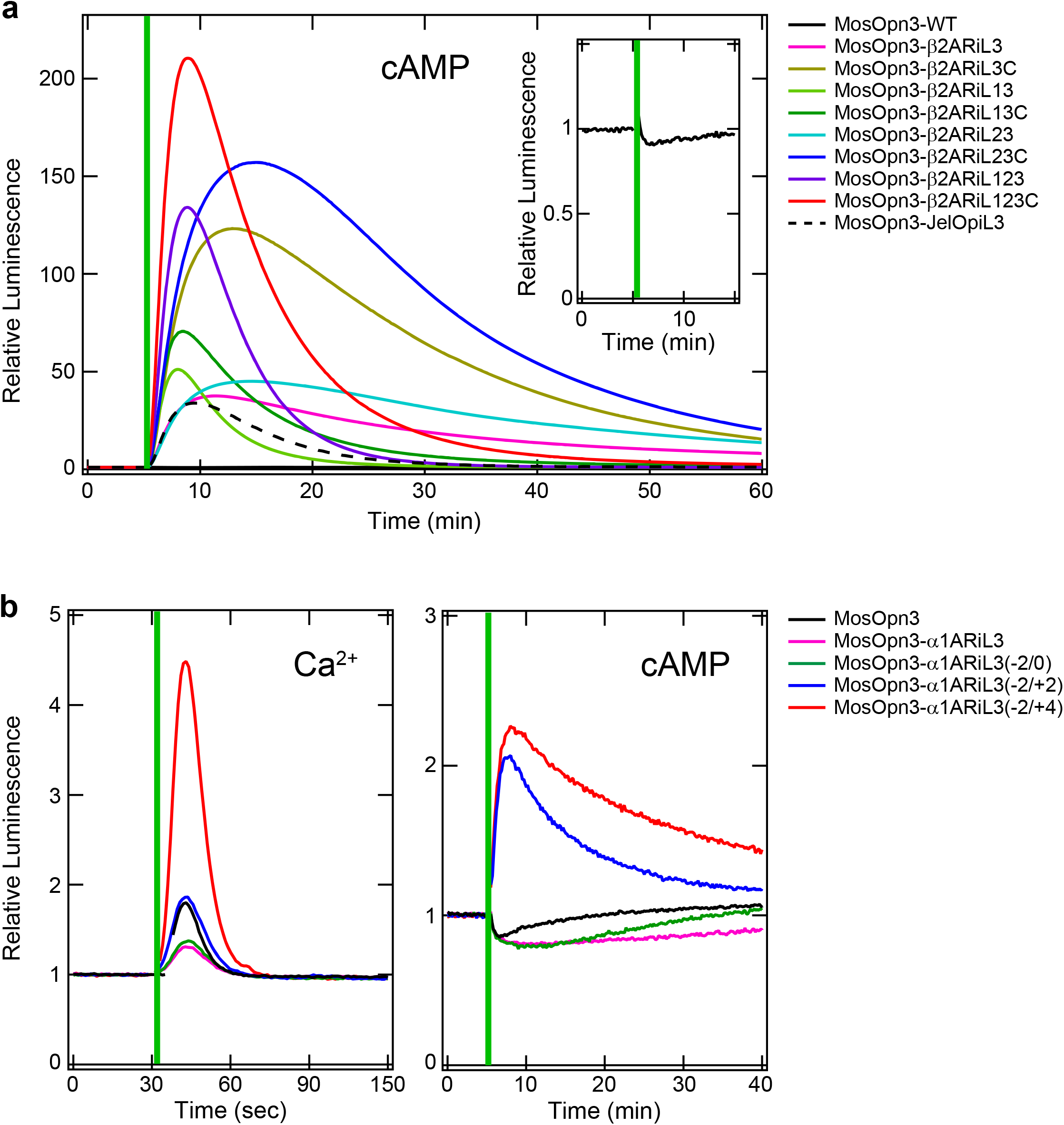
Engineering MosOpn3-based tools to activate Gs or Gq-type G protein. **a** GloSensor cAMP assay with HEK293 cells expressing various MosOpn3-ß2AR chimeras, showing light-induced increases of intracellular cAMP levels. MosOpn3 chimera containing the third cytoplasmic loop of jellyfish Gs-coupled opsin (MosOpn3-JelOpiL3) was also shown for a comparison. The light-induced cAMP decease in HEK293 cells expressing MosOpn3 WT was shown in inset. Vertical lines indicate light illumination for 5 sec. **b** Aequorin Ca^2+^ assay with HEK293 cells expressing MosOpn3-α1ARiL3 chimeras having varied length of third cytoplasmic loop region of α1AR, showing light-induced increases of intracellular Ca^2+^ levels (left). Light-induced changes of intracellular cAMP levels of HEK293 cells expressing the MosOpn3-α1AR chimeras were also shown (right). Note that other MosOpn3-α1AR chimeras did not show significant improvement of light-induced Ca^2+^ elevation. Vertical lines indicate light illumination for 1 sec (left), and 5 sec (right).

Regarding the regulation of Ca^2+^ level, aequorin luminescence-based calcium assay revealed that MosOpn3 WT originally exhibited a light-induced increase in cultured cells (Fig. 3b), although it also induced a decrease of cAMP level as described previously^22^ (Fig. 3a, b). Then we evaluated specific regulation of Ca^2+^ by MosOpn3-α1AR chimeras in cultured cells expressing each chimera. Contrary to the case of MosOpn3-ß2AR chimeras, only the MosOpn3-α1AR chimera having the third cytoplasmic region of α1AR (MosOpn3-α1ARiL3) exhibited a light-induced Ca^2+^ increase under the condition without 11-*cis* retinal addition, although the Ca^2+^ increase by the chimera is less than that by the WT and the chimera still exhibited cAMP decrease (Fig. 3b). To improve the specificity of the MosOpn3-α1ARiL3 for Ca^2+^ increase, we further engineered the chimera by fine-tuning of the third cytoplasmic region to be exchanged by shifting the donor/acceptor boundaries by two or four amino acids. In all possible combinations, we found that chimeras generated by reduction of two amino acids at the N-terminal and addition of four amino acids at C-terminal of the third cytoplasmic loop of α1AR (MosOpn3-α1ARiL3(−2/+4)) exhibited the highest functionality in light-induced Ca^2+^ elevation, which is ~2.5-fold higher than the case of WT without cAMP decrease. The light-induced cAMP increases observed in the case of MosOpn3-α1ARiL3(−2/+4) and MosOpn3-α1ARiL3(−2/+2) were presumably caused by Ca^2+^ elevation^52–54^. These results suggest the practical strategy for creating on-demand optogenetic tools based on MosOpn3 for manipulating intracellular cAMP and Ca2+ levels probably through optimizing G protein signaling preference.

### Parapinopsin for color-dependent manipulation of GPCR signaling

In cases of visible light-sensitive bistable opsins including MosOpn3, the dark (inactive) and photoproduct (active) states have largely overlapped absorption spectra in the visible light region. On the contrary, in the case of parapinopsin, the absorption spectra of UV-sensitive inactive state and visible light sensitive active state are distinct from each other, enabling to illuminate only the active state with visible light, which leads to complete regeneration of inactive state. To evaluate a contribution of the photoregeneration ability in optical manipulation of GPCR signaling *in vivo*, we focused on another *C. elegans* behaviors, coordinated movement regulated by acetylcholine. Using the promoter of unc-17, we introduced LamPP into cholinergic motor neurons, in which activation of Go-mediated signal transduction down-regulates acetylcholine releases to lead uncoordinated movement including coiling of *C. elegans*^55,56^ (Fig. 4a, b). Tg worms expressing LamPP in cholinergic motor neurons (LamPP-worms) were obtained with the same procedure as that for MosOpn3 worms. When we illuminated LamPP-worms fed 11-*cis* retinal-containing *E. coli* with violet light, the Tg worms stopped moving and coiled as expected, and upon subsequent green light illumination, the Tg worms restarted coordinated movement (Fig. 4c, d, Supplementary Movie 2). The violet light-induced coiling of LamPP-worms sustained for 30 min after the turn-off of light, and subsequent green light illumination restored the movement (Fig. 4e). The behavioral switching between coiling and moving by violet and green light stimuli occurred repeatedly (Supplementary Movie 2), demonstrating that the *C. elegans* behaviors are manipulated in a color-dependent manner using LamPP. In other words, the result suggested that introducing LamPP into cells containing Gi/o-mediated signaling could render physiologies color-dependency. Furthermore, LamPP-worms fed all-*trans* retinal under the red light (600 nm) exhibited violet-light-induced stop and green-light-induced recovery like the case of LamPP-worms fed 11-*cis* retinal (Fig. 4d). Since retinal hardly absorbs red light, the phenomena can be explained by the complete photoregeneration ability of LamPP; LamPP bound all-*trans* retinal to form the active state (LamPP* in Fig. 4b) directly and the active state completely converted to the 11-*cis* retinal-binding inactive state by red light absorption.

**Fig. 4.**
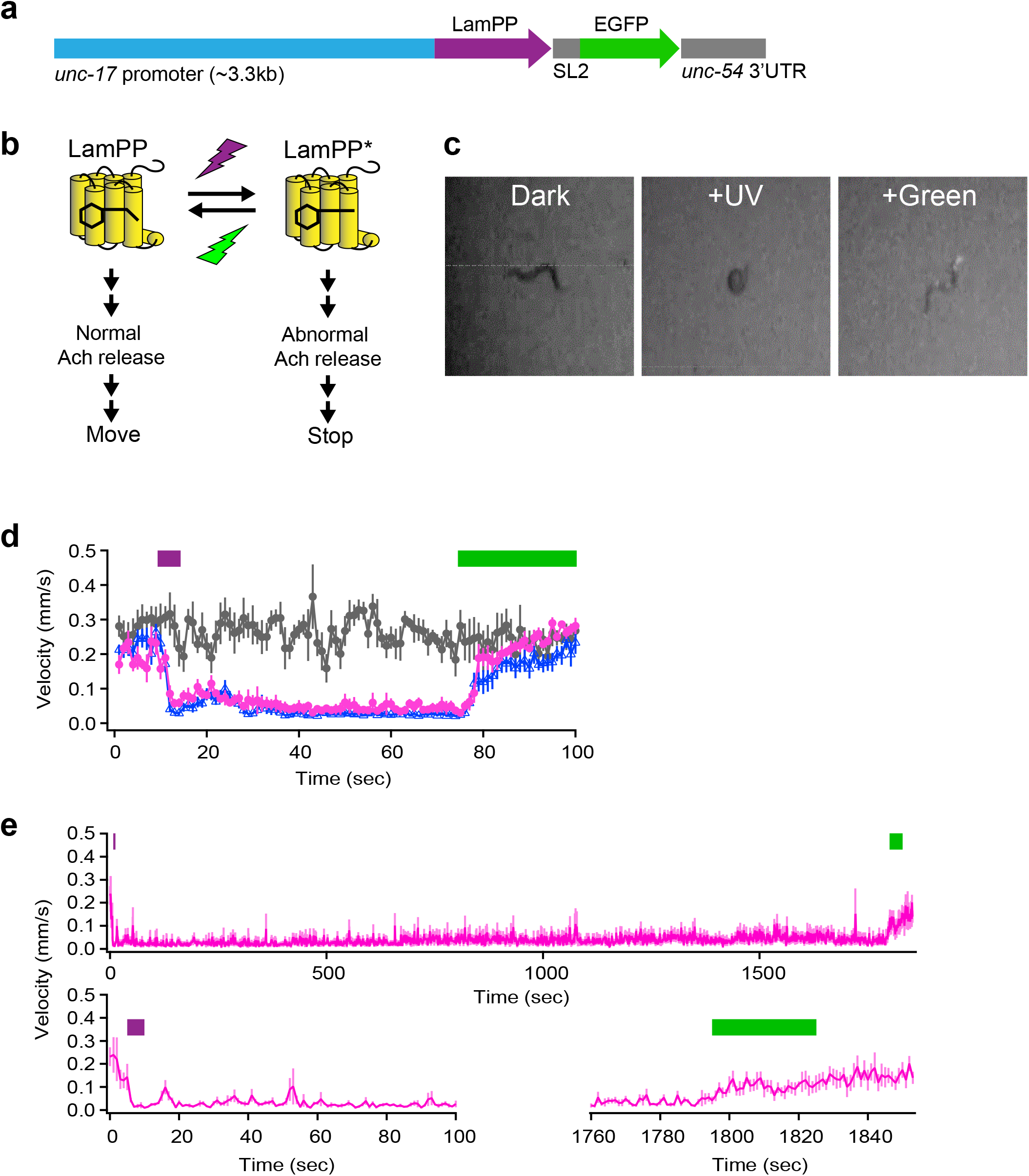
Color-dependent manipulation of Tg *C. elegans* expressing LamPP in cholinergic motor neurons. **a** The construction for introducing LamPP with EGFP under the *unc-17* promoter. **b** Schematic drawing of the relationship between status of LamPP and *C. elegans* behavior caused by violet and green light illumination. LamPP* indicates the active state of LamPP. **c** Snapshot images of the behavior of *C. elegans* expressing LamPP in cholinergic neurons in the dark, after violet illumination and subsequent green light illumination, showing moving forward, coiling and restart moving, respectively. See also Supplementary Movie 2. **d** Quantitative evaluation of violet light-induced coiling and subsequent green light-induced recovering of LamPP-worms (n=5). LamPP-worms fed *E. coli* without retinal (grey circles), with 11-*cis* retinal (magenta circles) in the dark and with all-trans retinal under the red light (blue circles) were illuminated with violet light for 5 sec (violet line), kept in the dark for 1min and illuminated with green light for 30 sec (green line). **e** Violet light-induced coiling of LamPP-worms fed 11-*cis* retinal sustained for 30 min after turn-off of the violet light. All worms exhibited immobility and/or coiling for 30 min. Upon green illumination, all worms restarted moving.

Again, we engineered LamPP, which originally activates Gi/o-type G protein, to activate Gs-type G protein to up- and down-regulate intracellular cAMP levels with violet and visible light absorption, respectively, which is in the opposite direction caused by the WT. We generated a series of LamPP chimeras, in which cytoplasmic region(s), including the third cytoplasmic region was replaced with those of ß2-adrenergic receptor as in the case of engineering Gs-coupled MosOpn3. Light-induced changes of cAMP level in cultured cells expressing each chimera were measured using the GloSensor cAMP assay. We found that only the cells expressing the chimera containing the third cytoplasmic loop alone (LamPP-ß2ARiL3) exhibited a significant violet light-induced increase of cAMP level, which in turn decreased to the basal level by subsequent green-light illumination (Fig. 5a). The color-dependent up- and down-regulations of cAMP levels occurred repeatedly. Notably, the cAMP increases induced by activation of the chimera were composed of two phases; upon violet-light illumination, cAMP levels rapidly increased to a higher level and immediately decreased to a lower level (acute phase) and the lower level sustained for more than 1 hour until green light illumination (chronic phase). Interestingly, when LamPP-ß2ARiL3-expressing cells were illuminated with blue light, the cAMP level was set at a level between levels caused by violet and green light illumination (Fig. 5b). Furthermore, the cAMP levels caused by blue light illumination were almost constant regardless of the levels just before illumination. The color dependency of sustained cAMP level could be explained in part by color dependent photoequilibrium of the inactive and active states of LamPP^17,35^. We also tested the performance of an additional LamPP chimera containing the third cytoplasmic region of the jellyfish Gs-coupled opsin, LamPP-JelOpiL3. The LamPP-JelOpiL3 exhibited much higher amplitude of violet light-induced increases compared with LamPP-ß2ARiL3, whereas the increased cAMP level gradually decreased (Supplementary Fig. 2), unlike the case of LamPP-ß2ARiL3. The increase and decrease of cAMP level by violet light and green light illumination also occurred repeatedly for LamPP-JelOpiL3. Collectively, these Gs-coupled LamPPs enable to manipulate Gs-mediated signal transduction in a color dependent manner, showing successful expansion of LamPP as tools for color-dependent manipulation of GPCR signaling.

**Fig. 5.**
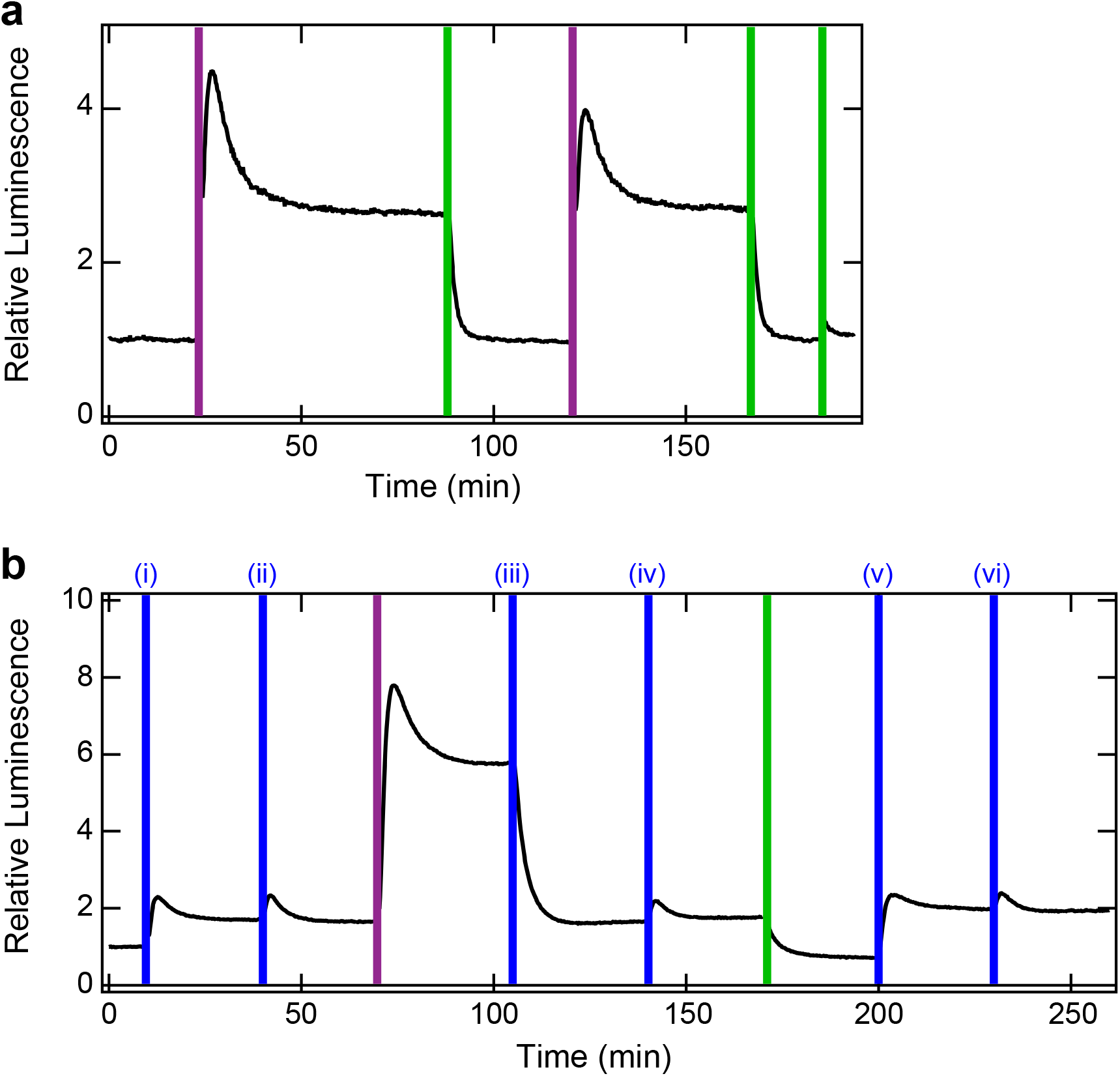
Color dependent manipulation of cAMP level by LamPP-ß2ARiL3 through the activation of Gs-type G protein measured by GloSensor cAMP assay. **a** Violet light illumination increased intracellular cAMP levels in HEK293 cells expressing LamPP-ß2ARiL3 and subsequent green light illumination decreased the cAMP level to the base level, which occurred repeatedly. **b** Blue light illumination generated a particular cAMP level. The cAMP levels generated by (i) 1st illumination, (ii) the subsequent illumination, (iii) the illumination after violet light illumination, (iv) the subsequent illumination, (v) the illumination after green light illumination and (vi) the subsequent illumination of blue light were almost constant, showing color-dependent cAMP clamping. Vertical lines indicate light illumination for 5 sec.

## Discussion

In this study, we demonstrated high performances of two bistable opsins, mosquito Opn3 (MosOpn3) and lamprey parapinopsin (LamPP) in manipulating GPCR signalings *in vivo*. Tg-worms expressing MosOpn3 in ASH neurons exhibited light-induced avoidance behaviors under the presence of 11-*cis* retinal and all-*trans* retinal with a similar sensitivity, which is comparable to the case of Tg-worms expressing bovine rhodopsin (BovRh) under the presence of 11-*cis* retinal (Fig. 1, 2). On the other hand, BovRh-worms did not exhibit light-induced avoidance behaviors under the presence of all-*trans* retinal as shown in the previous report examining the functionality of BovRh in other type of neurons of *C. elegans*, in which 9-cis retinal addition induced light-dependent neural responses but all-*trans* retinal addition did not^6^. Since all-*trans* retinal is basically present in every tissue, the perfomance of MosOpn3 under the presence of all-*trans* retinal comparable with that of BovRh under the 11-*cis* retinal suggests that MosOpn3 has a crucial advantage in optical manipulation of GPCR signaling *in vivo*. It is widely accepted that microbial rhodopsins such as ChR2 are powerful tools for use in optogenetics, especially in optical manipulation of neural activities^57–60^. One of the reasons for the popularity is that microbial rhodopsins form photosensitive pigments by binding all-*trans* retinal to function in every tissue, and the 11-*cis* retinal-requirement of bleaching opsins is the main reason for unpopularity of animal opsins. In terms of this point, the current study demonstrated that the availability of MosOpn3 could be equivalent to that of microbial rhodopsins. Moreover, the sensitivity of the avoidance response light-dependently induced by MosOpn3 in ASH neurons was revealed to be ~0.0002 mW/mm^2^ (white light) under the presence of all-*trans* retinal (Fig. 2c), which is ~7000 times higher than that induced by ChR2 in ASH neurons (~1.48 mW/mm^2^, blue light) reported in the previous study^61^. In fact, we generated Tg worms expressing ChR2 in ASH neurons and tested light-induced avoidance behaviors with the same protocol as in cases of MosOpn3 and BovRh but no reproducible avoidance response was observed by illuminating with ~0.8 mW/mm^2^ of white light, the highest light intensity of our light source. These data demonstrated that MosOpn3 could function as a light-sensitive switch to provide much higher photosensitivity to any tissue with much lower light-induced toxicity and/or temperature increase compared with the case of ChR2. There was a potential concern that the basal activity of 13-*cis* retinal-bearing MosOpn3 pigment might affect cellular conditions intrinsically *in vivo* because the activity of G protein by the 13-*cis* retinal-bearing pigment in the dark was higher than that of the 11-*cis* retinal-bearing pigment^22^. However, MosOpn3-worms fed all-*trans* retinal-containing *E. coli*, in which 13-*cis* retinal-bearing MosOpn3 pigments formed, normally behaved in the dark (Fig. 2). The data suggests that intracellular conditions in ASH neurons were somehow maintained to be normal, even with the partial G protein activation by 13-*cis* retinal-bearing MosOpn3 pigment. Then we established ß2AR (Gs-coupled receptor) and α1AR (Gq-coupled receptor) versions of MosOpn3 by replacing cytoplasmic region(s) with those of ß2AR and α1AR, respectively, to expand the applicability of MosOpn3 (Fig. 3). Analysis of series of MosOpn3 chimeras revealed that fine tuning of replaced regions improves the G protein activation efficiency and specificity (α1ARiL3) and alters the kinetics of chimeras (ß2AR), which could be a guide for making various MosOpn3 chimeras with other GPCRs of interest for appropriate purposes.

We also demonstrated color-dependent manipulation of GPCR signaling by LamPP to lead behavioral switching between coiled and moving of *C. elegans* by violet and green light stimuli, respectively (Fig. 4d, e). The switching is consistent with the molecular behavior of LamPP by UV and visible light absorption^17^, which suggests that the photoregeneration ability of LamPP contributes to the mode change of animal behaviors. Interestingly, we recently reported that in the zebrafish pineal photoreceptor cells, the parapinopsin alone generates color opponency; UV and orange lights induce hyperpolarization and depolarization, respectively, based on the photoequilibrium levels between inactive and active states of the parapinopsin^35^. The idea was also supported by current results by heterologous expressions of LamPP in *C. elegans* neurons, showing that introducing LamPP is sufficient for color-dependent behavioral switching. Furthermore, we demonstrated that LamPP-worms fed all-*trans* retinal under the red light (600 nm) responded to light stimuli (Fig. 4e), indicating that LamPP could function as a light-sensitive switch in every tissue with red light pre-illumination. The availability of LamPP is practically equivalent to MosOpn3 as well as microbial rhodopsins.

The color-dependent manipulation of Gi/o type G protein by the wild type LamPP has also been expanded to that of Gs type G protein by LamPP-ßARiL3 and LamPP-JelOpiL3, which enables color-dependent manipulation of intracellular cAMP levels in the reverse direction to the reactions by the wild type (Fig. 5). Interestingly, LamPP-ßARiL3 showed molecular properties for not only color-dependent switch-on and switch-off of GPCR signaling but also color-dependent and context-independent maintenance of intracellular cAMP level (Fig. 5). The results indicate that LamPP-ßARiL3 achieves clamping intracellular cAMP levels depending on color of light. We also emphasize that the cAMP levels were maintained in the DARK after a light flash, depending on its color, suggesting an alternative strategy for “light”-regulation of cAMP level. On the other hand, the “cAMP-clamping” was not achieved in the case of LamPP-JelOpiL3 (Supplementary Fig. 2). The clamped cAMP level could relate to shut off manners of the active state of LamPP-ßARiL3 based on unknown effects of the third cytoplasmic loop of ß2AR including phosphorylation and/or arrestin-binding, which provides a new insight into prolongation of GPCR signalings.

In this study, the availability of two bistable animal opsins has shown to be equivalent to that of microbial rhodopsins including ChR2 in terms of retinal requirement, which would accelerate research by optical manipulation of GPCR signaling. In manipulating GPCR signaling, chemical manipulation, namely chemogenetics using chemoreceptors such as DREADD (designer receptor exclusively activated by designer drugs) have been widely used^62–64^, which is in contrast to the situation of optogenetics with animal opsins, mainly bleaching opsins. The low popularity of optical manipulation of GPCR signaling was mainly due to the absence of effective and robust tools but the situation has been recently changing with bistable opsins, especially MosOpn3 and LamPP^28,36–38^, and advantages of them and their derivatives were systematically demonstrated based on their molecular properties in this paper. The main advantage of chemogenetics is that stimuli (chemicals) can be reliably delivered deep inside the body. On the other hand, temporally precise manipulation of GPCR signaling *in vivo* is basically unachievable by chemogenetics but achievable by optogenetics. In particular, temporally precise termination of G protein signaling as demonstrated by LamPP (Fig. 5) is a significant advantage of optogenetics, which enables controlling durations, intervals and numbers of the stimuli. Together with chemical manipulation, the optical manipulation with bistable animal opsins would greatly facilitate comprehensive and deeper understating of GPCR-based physiologies as well as GPCR signaling themselves.

## Materials and Methods

### Generating Tg *C. elegans*

The 2.4 kbp promoter sequence of *sra*-6^42^ was obtained from the *C. elegans* genomic DNA by PCR. The 3.3 kbp promoter sequence of *unc*-17 was a generous gift from Prof. Robert Lucas^56^. The vector backbone containing SL1 and GFP was obtained by digestion of the plasmid [pEM1 = flp-21::LoxPStopLoxP::npr-1 SL2 GFP] (Addgene plasmid # 24033)^65^ with NotI and KpnI. The *sra*-6 promoter linked with MosOpn3 cDNA or *unc*-17 promoter linked with LamPP cDNA was introduced into the vector. Each plasmid was co-injected into the wild type *C. elegans* strain N2 obtained from the Caenorhabditis Genetics Center with *pmyo-2::mCherry* as a selection marker. The wild and Tg strains (*psra-6::MosOpn3::GFP, punc-17::LamPP::GFP*) were cultured according to standard methods^66^.

## Behavioral experiments

Adult worms were used for behavioral experiments. Tg worms were fed *Escherichia coli* strain OP50 which are mixed with or without certain amounts of retinal at 24 h before experiments. The standard concentrations of 11-*cis* retinal or all-*trans* retinal in OP50 for our experiment were 0.38 mM or 0.67 mM, respectively, and diluted to 1/100 to 1/10000. Worms were monitored under infrared light illumination. White lights (I_0_ = ~0.8 mW/mm^2^) supplied by a 200 W metal-halide lamp (PhotoFluor II, 89 North) with or without ND filter(s) were applied as light stimuli for MosOpn3-worms. Narrow band violet (410 nm) and green (510 nm) LED lights were applied as light stimuli for LamPP. Red (600 nm) LED light was used as a background light while LamPP-worms were fed all-*trans* retinal. Avoidance response probabilities were calculated by the ratio or the number of responded worms to five worms examined.

### Generation of chimeric mutants of opsins

Chimeric mutants of MosOpn3 and LamPP were generated by replacing cytoplasmic regions of hamster ß2-adrenergic receptor (ß2AR) and human α1-adrenergic receptor (α1AR) by combining DNA fragments of each region with PCR. The cytoplasmic regions were determined according to previous reports^5,50^ and applied to MosOpn3 and LamPP based on the alignment including bovine rhodopsin.

### Bioluminescent reporter assays for Ca^2+^ and cAMP

The intracellular cAMP and Ca^2+^ levels in opsin-expressing HEK293S cells were measured using the GloSensor cAMP assay and the aequorin assay, respectively, as described previously^22,26,67,68^. The pGloSensor-22F cAMP plasmid (Promega) was used for the GloSensor cAMP assay. The wild type aequorin obtained by introducing two reverse mutations into the plasmid [pcDNA3.1+/mit-2mutAEQ] (Addgene #45539)^69^ was used for the aequorin assay. A broad band green LED light and narrow band violet (410 nm) and blue (430 nm) LED lights were applied for 5 sec in the GloSensor cAMP assay and for 1 sec in the aequorin assay as light stimuli.

## Acknowledgements

We thank Robert J. Lucas (The University of Manchester) for providing the promoter of unc-17. This work was supported in part by grants-in-aid for Scientific Research from the Japanese Ministry of Education, Science, Sports and Culture 18H02482, 20K21433, 21H00435 (to M.K.), 15H05777, 20K21434 (to A.T.); Japan Science and Technology Agency (JST) Precursory Research for Embryonic Science and Technology (PRESTO) Grant JPMJPR13A2 (to M.K.) and JST Core Research for Evolutional Science and Technology (CREST) Grant JPMJCR1753 (to A.T.).

## Author Contributions

M.K. and A.T. conceived and designed the study. M.K., B.S., T.N., L.S., S.W., S.K., and E.K.-N. carried out the experiments and analysed the data. M.K. and A.T. wrote and edited the paper.

