## Supplementary Figure1-2 for "High-performance GPCR optogenetics based on molecular properties of animal opsins, MosOpn3 and LamPP"

### Supplementary Figures

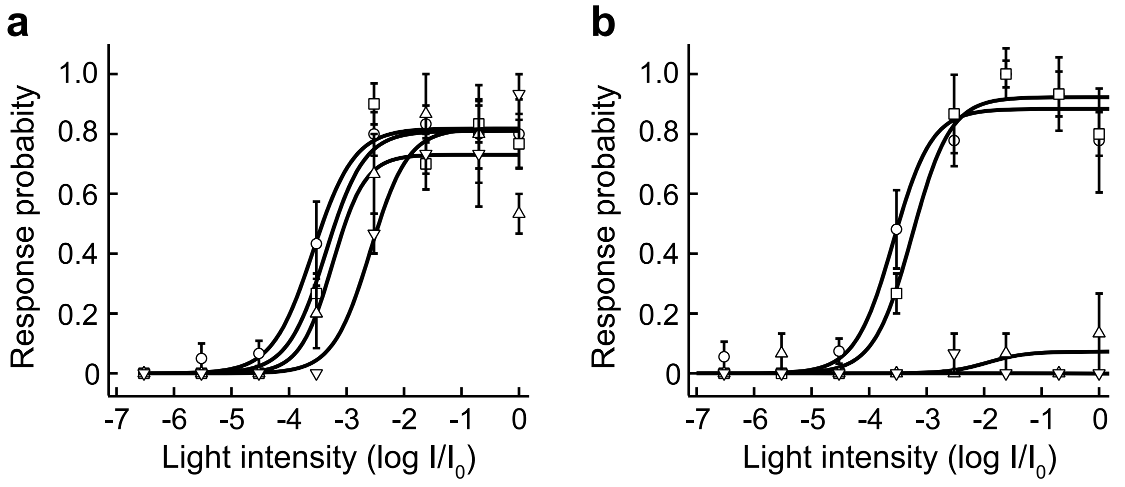


Supplementary Fig. 1 Koyanagi et al.

**Supplementary Fig. 1. Light intensity-response probability relationships of MosOpn3- and BovRh-worms with various amounts of 11-*cis* retinal.**

**a** MosOpn3/11-worms. **b** BovRh/11-worms. The amount of 11-cis retinal added to worms were 1/1 (circles), 1/100 (squares), 1/1000 (triangles) or 1/10000 (inverted triangles) of the standard. n=3~5.

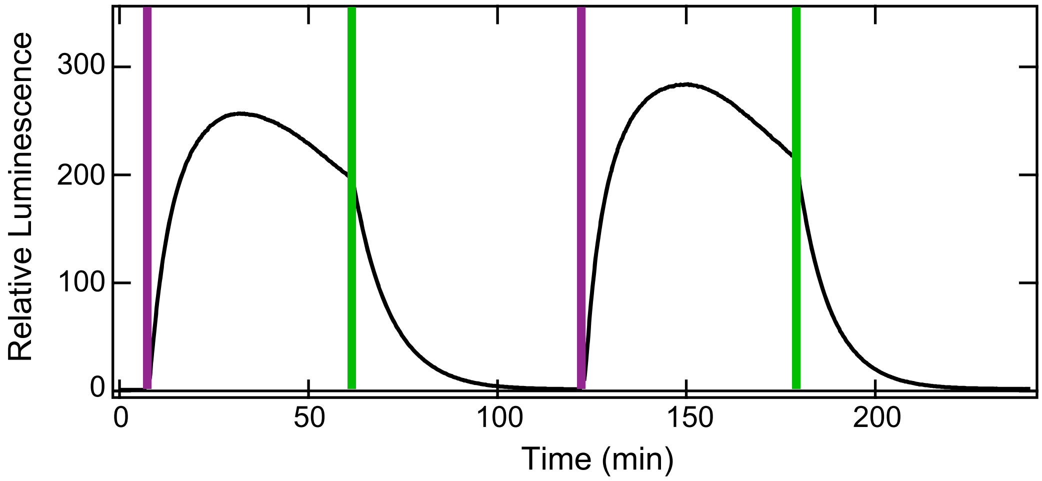


Supplementary Fig. 2 Koyanagi et al.

**Supplementary Fig. 2. Color-dependent manipulation of cAMP level by LamPP-JelOpiL3 through the activation of Gs-type G protein measured by GloSensor cAMP assay.**

Violet light illumination increased intracellular cAMP levels in HEK293 cells expressing LamPP-JelOpiL3 and subsequent green light illumination decreased the cAMP level to the basal level, which occurred repeatedly.

### Supplementary Movie legends

**Supplementary Movie 1. Movies showing typical light-induced avoidance behaviors of Tg *C. elegans* expressing MosOpn3 or BovRh in ASH neurons.**

**a** MosOpn3/11-worm. **b** MosOpn3/NoRet-worm. **c** MosOpn3/AT-worm. **d** BovRh/11-worm. **e** BovRh/NoRet-worm. **f** BovRh/AT-worm. Timings of light illumination were indicated by emergence of white frames.

**Supplementary Movie 2 The movie showing color-dependent manipulation of behaviors of Tg *C. elegans* expressing LamPP in cholinergic motor neurons.**

A LamPP-worm was moving before illumination, stopped moving and coiled upon violet light illumination, and restarted movement upon green light illumination, which occurred repeatedly. Timings of violet and green light illumination were indicated by emergence of violet and green frames, respectively. Signals derived from mCherry were also detected by green light illumination.
